# Genetic Landscape of Gullah African Americans

**DOI:** 10.1101/2020.10.12.336347

**Authors:** Kip D. Zimmerman, Theodore G. Schurr, Wei-Min Chen, Uma Nayak, Josyf C. Mychaleckyj, Queen Quet, Lee H. Moultrie, Jasmin Divers, Keith L. Keene, Diane L. Kamen, Gary S. Gilkeson, Kelly J. Hunt, Ida J. Spruill, Jyotika K. Fernandes, Melinda C. Aldrich, David Reich, W. Timothy Garvey, Carl D. Langefeld, Michèle M. Sale, Paula S. Ramos

## Abstract

**Objectives:** Gullah African Americans are descendants of formerly enslaved Africans living in the Sea Islands along the coast of the southeastern U.S., from North Carolina to Florida. Their relatively high numbers and geographic isolation were conducive to the development and preservation of a unique culture that retains deep African features. Although historical evidence supports a West and Central African ancestry for the Gullah, linguistic and cultural evidence of a connection to Sierra Leone has led to the suggestion of this country/region as their ancestral home. This study sought to elucidate the genetic structure and ancestry of the Gullah.

**Materials and Methods:** We leveraged whole-genome genotype data from Gullah, African Americans from Jackson, Mississippi, Sierra Leone Africans, and population reference panels from Africa and Europe, to infer population structure, ancestry proportions, and global estimates of admixture.

**Results:** Relative to southeastern non-Gullah African Americans, the Gullah exhibit higher mean African ancestry, lower European admixture, a similarly small Native American contribution, and stronger male-biased European admixture. A slightly tighter bottleneck in the Gullah 13 generations ago suggests a largely shared demographic history with non-Gullah African Americans. Despite a slightly higher relatedness to Sierra Leone, our data demonstrate that the Gullah are genetically related to many West African populations.

**Discussion:** This study confirms that subtle differences in African American population structure exist at finer regional levels. Such observations can help to inform medical genetics research in African Americans, and guide the interpretation of genetic data used by African Americans seeking to explore ancestral identities.

**Research Highlights:**

- Using genomic data, we show that the Gullah have lower European and higher West African genomic background compared to non-Gullah African Americans, confirming their diverse African ancestry and rejecting a model that asserts a predominant Sierra Leone origin.
- Our data reveal a largely shared demographic history with southeastern non-Gullah African Americans, but also subtle differences related to high African genetic ancestry due to isolation in the Sea Islands.

## INTRODUCTION

Knowledge about the genetic background of a population, including regional differences in ancestry, structure, and variation, is not only critical for medical and population genetic studies (1), but can also illuminate questions of social and cultural relevance. Multiple studies show wide variability in the levels of African ancestry in African American individuals in the United States (U.S.) at both state and regional levels (2-8). This variation in African ancestry underscores the importance of properly accounting for ancestry in historical and biomedical studies of diasporic populations (9, 10).

While regional patterns in ancestry proportions in African Americans in the U.S. are broadly understood, fine-scale characterization of the ancestral diversity of discrete groups is lacking. In addition, as a result of the trans-Atlantic slave trade, African Americans were robbed of their African heritage and left with limited information about ancestors and homelands (11). The longing for identity and belonging leads many African Americans to actively draw together and evaluate various sources of genealogical information (historical, social and genetic) in order to weave together ancestry narratives (12). Elucidating the African ancestries of African Americans can enrich the lives of African Americans by helping them to enrich a sense of identity, make connections to ancestral homelands, and ultimately foster reconciliation in the wake of emancipation (13).

The Gullah are a culturally distinctive group of African Americans from the coastal Sea Islands of North Carolina, South Carolina, Georgia, and Florida. On many plantations of the coastal Sea Islands, Africans vastly outnumbered Europeans. Relative isolation fostered the development of a unique culture in which many African influences were preserved, including language, folktales, religious beliefs, food preferences, music, dance, arts, and crafts (14, 15).

Marked by unique intonation and rhythm as well as syntax and lexicon, the Gullah language is hardly intelligible to the outsider, and remains the most characteristic feature of the sea islanders. The origin of this unique Creole language, like the origin of the Gullah people, is still debated. One hypothesis proposes that they descend from Krio ancestors originating in Sierra Leone (16, 17), a view supported by the fact that contemporary sea islanders can understand the Krio of Sierra Leone and vice versa. This striking linguistic resemblance, coupled with multiple cultural links (e.g., rice growing techniques, quilts, songs, stories), has led to a commonly held folk history that the Gullah are descendants of enslaved Africans from the African Rice Coast (16), the traditional rice-growing region stretching south from Senegal to Sierra Leone and Liberia (Supporting Figure S1).

However, several historical accounts support a diverse African ancestry of the Gullah (18-20). The recorded legal slave trade into Charleston, South Carolina, documents approximately 39% of enslaved Africans as originating from West Central Africa (present day Angola, Congo, and part of Gabon), 20% from Senegambia (present day Senegal and Gambia), 17% from the Windward Coast (present day Ivory Coast and Liberia), 13% from the Gold Coast (present day Ghana), 6% from Sierra Leone (present day Sierra Leone and Guinea), and 5% from the Bights of Benin and Biafra (Togo, Benin, Nigeria, Cameroon, and part of Gabon) (Supporting Figure S1) (18). In addition to cultural links (e.g., religious beliefs, arts and crafts), words and syntax support a larger role of West-Central Africa, the Gold Coast, and adjacent Nigeria in forming the Gullah language (21). While it has been proposed that the Congo-Angola area had an early cultural dominance with artifacts, lexicon and beliefs, the complexities of the Bantu grammar probably prevented its adoption in the Sea Islands. Senegambia, Sierra Leone, and the Windward Coast down through the Bight of Biafra contributed most in the latter half of the 18^th^ century, when half of all slaves imported into Charleston arrived, adding more words, grammar, and even whole stories (18). Thus, it is likely that, instead of Gullah deriving directly from Krio, both languages share a close common origin (18).

In this context, it is currently unknown how significant of a genetic trace that Sierra Leone ancestors might have left in present-day Gullah African Americans. Early genetic studies of autosomal, mtDNA, and Y-chromosome markers, indicated that the Gullah had high African ancestry (14, 22) and a lower genetic distance to populations from Sierra Leone compared with African Americans from urban areas (23, 24). Sierra Leone officially has sixteen ethnic groups, each with its own language and customs. The largest native ethnic groups include the Temne (35%) in northern Sierra Leone and areas around the capital, who arrived during the 11^th^ and 12^th^ centuries upon the fall of the Jalunkandu Empire in present day Republic of Guinea (25-27). The Mende (31%), who live mostly in the Southeast and the Kono District, originated in a region near western Sudan, but migrated from the inland to the coast between the 2^nd^ and 16^th^ centuries to trade woven cloths for salt (25-27). The Limba (8%) are native to the savannah-woodland region in northern Sierra Leone (25-27). By contrast, the Fula (7%) are descendants of Fulani migrant from Guinea who settled in Sierra Leone during the 17^th^ and 18^th^ centuries (25-27). Likewise, the Kono (5%) and the Mandingo (2%) are descendants from Guinea migrants (27) (Supporting Figure S2).

The Creole (2%) are descendants of freed African slaves from America who settled in Sierra Leone after 1787, and as such have multiple African origins (25). Following the American Revolutionary War (or War of Independence, 1775-1783), the British government freed Africans who served in the British armed forces and resettled them in Granville Town, the predecessor of Freetown and the present capital of Sierra Leone. Maroons, runaway enslaved Africans from the West Indies who formed independent settlements on different islands, were also resettled in Freetown, as were over 50,000 “recaptives” brought there by the British navy (28). The subsequent generations born in Sierra Leone were called Krio, Kriole, or Creole. In addition, individuals from other Sierra Leone ethnic groups joined the Creole communities, thereby promoting a fusion of African and Western cultures (29). The Krio language unites the different ethnic groups for trade and interactions with each other (30). The present national boundary was only fixed in 1896, prior to which people moved freely through the coastal country, making their own settlements, and fixing their own boundaries between themselves and their neighbors (25).

This study sought to elucidate the population structure of the Gullah and their relationship to contemporary Sierra Leone ethnic groups and other West African populations using genome-wide genotype data. African Americans from the Jackson Heart Study (JHS) (31, 32) recruited in Jackson, Mississippi, were also included to provide a comparison with a less geographically isolated, but regionally close, Southeastern U.S. African American sample. The results of this analysis support the multi-African ancestry and reduced European admixture of the Gullah compared to other U.S. African American populations. These results are consistent with historical data (18), which indicate that the Gullah are a mixture of numerous people from different genetic, ethnic, and linguistic currents who formed their own culture and language. Identifying the diverse people who played a role in shaping the Gullah has implications for all African Americans, and for the legacy of the African diaspora everywhere (18).

## SUBJECTS AND METHODS

The participants and methods employed in this study are described in greater detail in Supporting Information Subjects and Methods.

### Community engagement

This study was conducted with the approval and cooperation of the Sea Island Families Project (SIFP) Citizen Advisory Committee, representing a generative partnership between academic researchers and Gullah African Americans in rural South Carolina (33).

### Sample collection and SNP data generation

Self-identified Gullah African American subjects and their parents were born and raised in the Sea Islands region of South Carolina (along the coastal border and 30 miles inland). After obtaining informed consent, 5 mL blood samples were drawn from all participants, and DNA was extracted from them using a standardized DNA isolation kit (Gentra Systems, Minneapolis, MN). Sample collection and processing for the African American subjects from the Jackson Heart Study (JHS) (31,32), Sierra Leone (15), and Native American Mixtec subjects (34) have been previously described. DNAs from the Gullah and Sierra Leone African subjects were genotyped using the Affymetrix Genome-Wide Human SNP Array 6.0. After quality control (QC) procedures, 883 unrelated Gullah African Americans, 381 unrelated Sierra Leone Africans, 7 Mixtecs, and 1,322 unrelated JHS African Americans were retained for analyses (Supporting Table S1). We also included 125 HGDP (35) and 386 HapMap III (release 3) (36) individuals. The geographic distribution and linguistic affiliation of the African populations used in this study are shown in Supporting Table S2 and Figure 1A.

### Data merging and SNP trimming

PLINK v1.9 (37) was used to combine our Affymetrix 6.0 data and the HapMap III and HGDP data. After merging samples, 136,878 common variant SNPs met QC thresholds. For methods that required a set of linkage disequilibrium (LD)-pruned SNPs (see below), we further removed SNPs with an r^2^ > 0.1, leaving 64,303 SNPs for analysis. For more specific analyses, such as the IBD and IBDNe analyses, identical merging and filtering methods were used to combine the Affymetrix 6.0 data with the HapMap III data. This step resulted in a combined set of 579,854 filtered (but not pruned) SNPs for the Gullah and 615,569 for the non-Gullah (JHS) African Americans. For the qpAdm analyses, the Gullah and JHS data were combined with the HapMap III data and the Mixtec samples. This dataset consisted of 544,517 filtered (but not pruned) SNPs.

### Principal component analysis for inference of population structure

Principal component analysis (PCA) as implemented in EIGENSOFT v6.0.1 (38) was computed using the set of LD-pruned 64,303 SNPs described above.

### Global estimates of admixture

Unsupervised clustering as implemented in ADMIXTURE v1.3.0 (39) was used to estimate global genetic ancestry of the European, African, and African American individuals, assuming 2 through 8 ancestral genetic clusters (*k*=2 through *k*=8) to determine the optimal number of ancestral reference groups. Five clusters (i.e., *k*=5) gave the lowest cross-validation error (Supporting Figure S3). To help order the populations according to their genetic similarities, we used an average linkage hierarchical cluster analysis based on the means of each of the five ancestral populations computed by ADMIXTURE and inter-population similarity matrix of Euclidean distances. The results are presented in a dendrogram (Supporting Figure S4).

### Inference of ancestry proportions

To estimate ancestry proportions on autosomes and the X-chromosome for both the Gullah and non-Gullah (JHS) African-Americans, we used qpAdm (40). The HapMap LWK, MKK, CHB, JPT, GIH, and TSI populations were used as outgroups, and CEU, YRI, and Mixtecs (34) were used as proxies for source populations for European, African, and Native American ancestry, respectively.

### Detection of genomic segments shared identical-by-descent (IBD) between African American groups and estimation of effective population sizes

We used GERMLINE v1.5.1 (41) to infer IBD tracts of length 18 cM or longer that were shared between Gullah and non-Gullah (JHS) African American individuals. Ancestry-specific IBDNe (42) was used to estimate the effective population size for recent generations within each African American subgroup.

### Genetic diversity and population differentiation

Heterozygosity (HET) and inbreeding coefficients *(F)* were calculated using genotypic data on 273 healthy Gullah African Americans and 381 Sierra Leone individuals. Using genotype data from healthy Gullah, JHS African Americans, Sierra Leone, YRI and CEU samples, we computed the Weir and Cockerham’s (1984) F_ST_ (43) as implemented in VCFtools v0.1.13 (44).

## RESULTS AND DISCUSSION

### Genetic structure of Sierra Leone Africans

Given the general folk belief that Sierra Leone was the ancestral source of most Gullah African Americans, we first sought to characterize the population structure of African ethnic groups from this region. We combined genotype data from Sierra Leone Africans with Africans from the HGDP and HapMap III studies (Supporting Table S1), and inferred patterns of population structure and individual ancestry by principal component analysis (PCA) (Figure 1) and ADMIXTURE (39) (Figure 2). Consistent with previous reports (see the review by (45)), PCA distinguished geographic and linguistic African subpopulations, separating a combination of geographic groups and speakers of the four major language families (Afro-Asiatic, Nilo-Saharan, Niger-Kordofanian, and Khoisan) (Supporting Table S2).

**Figure 1.**
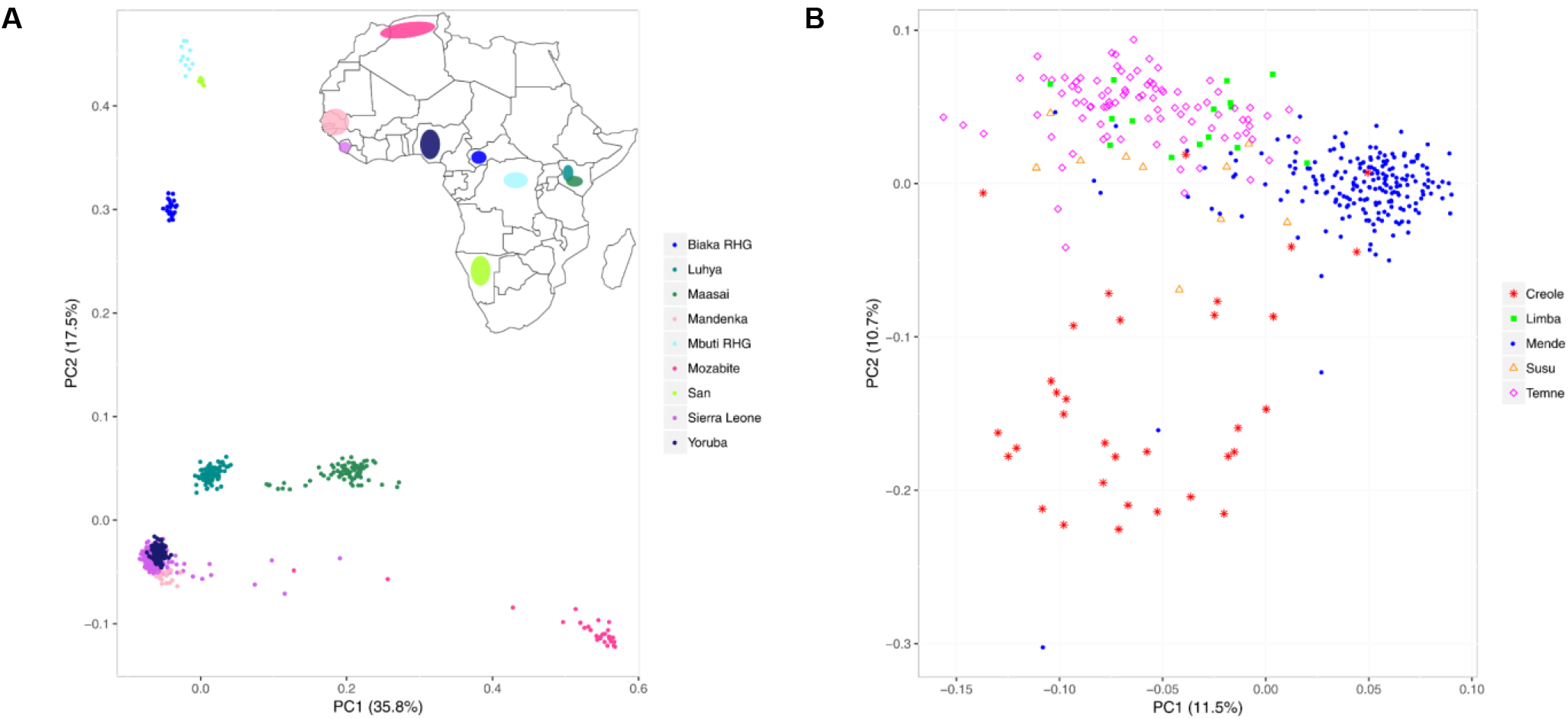
Principal component analysis of all African samples. Principal component analysis (PCA) (EIGENSOFT) was applied to HGDP and HapMap III African and Sierra Leonean populations. (A) PCA showing the Mozabite cluster (along PC1), the West, Eastern and Southern, and Middle and South Western subpopulations clusters (along PC2). Insert shows approximate locations of sampled populations in Africa. (B) PCA of Sierra Leone ethnic groups with n>10 showing the Mende, Creole and Temne forming relatively different, but overlapping, clusters.

As shown in Figure 1A, the first principal component (PC1) differentiated the Mozabites of North Africa from all other African populations. A few Mozabite and Sierra Leone individuals formed a geographical gradient, reflecting different levels of African and West-Eurasian-related admixture (Supporting Figure S5) (46). A slight partitioning of the East African groups also occurred, with the Maasai (Kenya) being separated from the Luhya (Kenya). PC2 further separated four main groups, with West Africans forming one cluster and the East African groups clustering together (Luhya and Maasai). The Biaka rainforest hunter-gatherers (Central African Republic) formed an individual cluster, and the Mbuti rainforest hunter-gatherers (Democratic Republic of the Congo) and San (Namibia) formed a more distant cluster. The relationship between population structure and geographic and linguistic factors is supported by the results from the global ancestry estimates (Figure 2 and Supporting Figure S4). ADMIXTURE (39) (Figure 2) showed the highest West African-associated ancestry in the Mandenka of Senegal and Sierra Leone ethnic groups (red), the highest East African-associated ancestry in the Maasai and Luhya (yellow), and the highest Central and South African-associated ancestry in the Mbuti and San, respectively (orange).

PCA was applied to data from Sierra Leone ethnic groups to further discriminate any potential clusters of genetic variation within the region (Figure 1B). Our results showed that the most populous ethnic groups (Mende and Temne) form relatively different, but overlapping clusters. The Limba clustered among the Temne, which was unexpected given their distinct, unrelated language, and an early analysis of mtDNA genetic diversity purporting that the Limba could be distinguished from the Mende, Temne, and Loko groups (15). The Limba are indigenous to Sierra Leone, and their dialects are largely unrelated to the other languages in the region. The Krio or Creole formed a relatively distinct cluster along PC2. This pattern of population structure was broadly consistent with individual ancestry estimates (Figure 2 and Supporting Figure S4), where the Temne and Mende showed similar ancestry proportions, and the Creole appeared more variable in their African ancestry than other groups (Figure 2 and Supporting Figure S4). The Creole were also slightly more similar to the Yoruba, while other Sierra Leone ethnic groups showed more genetic similarity to the Mandenka (Figure 3 and Supporting Figure S4).

The genetic composition of the Creole was intermediate between that of other Sierra Leone ethnic groups and the Yoruba, with ancestral diversity similar to that of the Gullah African Americans. This finding suggested that they were likely the descendants of individuals from various parts of Africa, including Sierra Leone and beyond, as well as African descended individuals with European admixture. This finding is also consistent with their demographic history, which suggests the Creole descended from freed enslaved Africans (25, 28) who mixed with other ethnic groups (29). In summary, analysis of the African samples in this study showed that, despite the similarity among African Rice Coast populations (Mandenka and Sierra Leone ethnic groups), most of the recognized ethnic groups exhibit considerable genetic diversity.

**Figure 2.**
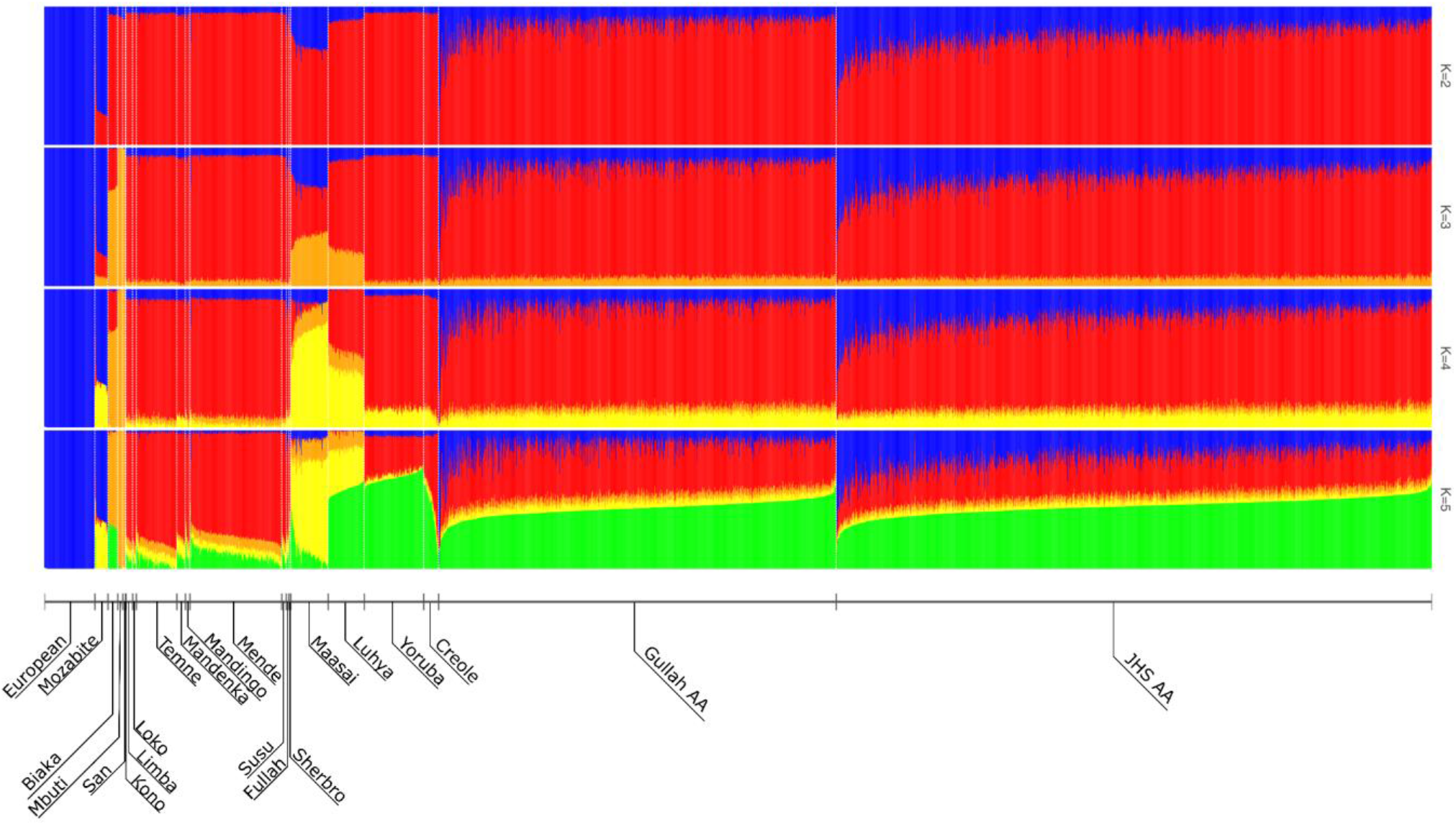
Ancestry estimates for European, African, and African American populations. ADMIXTURE analysis in Europeans, Africans (including Sierra Leone ethnic groups) and African Americans, assuming two through five ancestral genetic clusters (*k*=2 through *k*=5). The *k*=5 setting has the lowest cross-validation error of *k*=2-8. Populations were ordered via hierarchical cluster analysis. The plot shows each individual as a thin vertical column colored in proportion to their estimated ancestry from one particular population. The initial distinction is between Europeans (blue) and Africans (other colors). Within Africans, red indicates a West African (aka African Rice Coast) ancestry (highest in the Mandenka of Senegal and Sierra Leone ethnic groups), orange a Central and South African ancestry (highest in the San of Namibia and Mbuti of Democratic Republic of the Congo), yellow an East African ancestry (highest in Maasai and Luhya of Kenya), and green a West-Central African ancestry (aka Bight of Benin) ancestry (highest in the Yoruba of Nigeria).

### African ancestry estimates of Gullah African Americans

To characterize the global patterns of ancestry and population structure of Gullah African Americans, we used qpAdm (40, 47), ADMIXTURE (39), and PCA, with data from Gullah living in South Carolina, and for comparison with a regionally close group, from non-Gullah Southeast African Americans from the Jackson Heart Study (JHS). This analysis confirmed previous genetic reports of autosomal, mtDNA, and Y-chromosome markers, i.e., that Gullah African Americans had lower European admixture and higher African ancestry than other African American populations in the USA (14, 22-24). The higher average proportion of African ancestry in the Gullah was evident from autosomal global ancestry inference from qpAdm (40, 47), with the average African contribution to the Gullah African Americans being 90.7% compared with 82.2% in JHS African Americans (Table 1). Among studies of African ancestry in different U.S. regions (2-6, 8), the high African ancestry proportion seen in the Gullah was nearly matched in a U.S. cohort of African Americans sampled within rural Southeast U.S. (89% in Florida and 88% in South Carolina) (3). Thus, the Gullah show the highest average African ancestry proportion of any U.S. African American group studied to date.

**Table 1.** Estimates of African, European, and Native American ancestry in two African American cohorts (% with 95% confidence intervals)

|  | Ancestry |  |  |
| --- | --- | --- | --- |
|  | African | European | Native American |
| **JHS African Americans (n=1322)** |  |  |  |
| Autosomes (P-value = 0.00) | 82.2% [82.1 - 82.3] | 16.3% [16.2 - 16.4] | 1.4% [1.3 - 1.5] |
| X-chromosome (P-value = 0.14) | 86.5% [86.0 - 87.0] | 11.8% [11.2 - 12.2] | 1.7% [1.0 - 2.4] |
| Difference in mean X and autosomal ancestry | +4.3% | -4.5% | +0.3% |
| Estimated proportion of ancestry from males | 42.15% | 91.41% | 17.86% |
| **Gullah African Americans (n=883)** |  |  |  |
| Autosomes (P-value = 0.00) | 90.7% [90.6 - 90.8] | 8.0% [7.9 - 8.1] | 1.3% [1.2 - 1.4] |
| X-chromosome (P-value = 0.26) | 93.4% [92.9 - 93.9] | 4.3% [3.7 - 4.9] | 2.2% [1.1 - 3.1] |
| Difference in mean X and autosomal ancestry | +2.7% | -3.7% | +0.9% |
| Estimated proportion of ancestry from males | 45.53% | 119.38% | -53.85% |

Mean estimates (95% confidence intervals) of African, European, and Native American ancestry are shown. qpAdm was used to estimate proportions of European, African, and Native American ancestry, and qpAdm rank P-values are listed for both cohorts for each of the autosomes and chromosome X analyses. The mean X-chromosomal ancestry minus the mean autosomal ancestry (%) is listed. To aid with interpretation, the proportion of ancestry that comes from males was estimated under a simple model in which, in a population with equally many females and males, the mean X-chromosomal admixture fraction is a linear combination of female and male admixture parameters, with coefficients 2/3 and 1/3, respectively. The proportion of European ancestry across all chromosomes is higher in the Jackson Heart Study (JHS) relative to the Gullah African Americans, but the proportion of European ancestry that comes from males is higher in the Gullah African Americans.

In parallel with the higher average African ancestry, the European ancestry estimate in the Gullah was the lowest reported for African Americans in the U.S., while that for the JHS African Americans was similar to estimates in other groups (e.g., as low as 14% and 15%) (2, 3, 5) (Table 1). Consistent with the ancestry estimates, genome-wide admixture analyses showed that, despite the highly variable levels of European and West African ancestry in all African American groups, the Gullah had a lower average level of European admixture than JHS African Americans (Figure 2). The lower European contribution in the Gullah corroborates known differences in ancestry proportions among African Americans in different U.S. states (2-6, 8), and confirms that subtle differences in African American population structure can exist at finer regional levels.

The higher mean level of African ancestry in the Gullah is likely the result of the historically higher proportion of African Americans living in the Sea Islands of South Carolina since the early 1700s. African descendants comprised the majority of the population of South Carolina until the Great Migration to northern industrial cities in 1910 (48). As the demand for enslaved Africans to work in the rice fields, and later in the cultivation of indigo and cotton, was very high through the 18^th^ century and into the 19^th^ century, there was a large influx of Africans into South Carolina and Georgia since the beginning of the colonies (22). In the first federal census of 1790, enslaved Africans comprised 18% of the nation’s total population, but ranged from 47–93% in several coastal areas of South Carolina, including the port of Charleston, the center of American slavery (22). During that time, the Beaufort and Charleston Districts had 76% enslaved Africans, and the parish that included the Sea Islands was comprised of 93% enslaved African (49). Additionally, in the Sea Islands, all of the plantation owners who could leave the plantations from late May to late June would do so to avoid risk of contracting malaria. The absence of planters during this period gave the African descended individuals more autonomy in developing their own culture (18).

Later, in 1860, when the enslaved African population in the US declined to 13%, that of South Carolina had risen to 57%. The approximately equal number of male and female slaves in these districts by 1810 suggests that the increase was due to reproduction of local populations rather than the importation of individuals from Africa. The increase in the number of Africans, their concentration in rural areas, the severity of slave codes, and the social alienation of Africans from Europeans, produced isolation and a bond of brotherhood among the 18^th^ century Gullah people. These conditions provided an ideal context for creolization and the development of distinctive cultural attributes that continued into the 19^th^ century and beyond (18).

We also investigated whether the higher mean level of African ancestry in the Gullah could be a direct effect of a potentially higher proportion of African Americans currently living in this region. Currently, in its area of residence, the Gullah community sampled for this study encompasses 25-49% of the population (50), which is similar to the proportion of African Americans in the tri-county area sampled for the Jackson Heart Study (50). The slightly higher proportion of African Americans in Mississippi (37% vs. 28% in South Carolina) (50) suggests that the reported differences in population structure were not simply the result of current differential population proportions. Given the similar proportions of African Americans living in the Mississippi and South Carolina counties sampled for this study, the higher mean level of African ancestry in the Gullah is not likely an effect of the current population proportions, but rather reflects the historically higher proportion of African Americans living in the Sea Islands of South Carolina since the early 1700s until the mid 1900s.

Finally, since African Americans living in rural areas have a higher average African ancestry than those living in urban areas (3), we further considered the effects of sampling on ancestry proportions. The Gullah are an intrinsically rural community, while JHS participants represent urban dwellers. Baharian and colleagues (3) report that, for both African Americans sampled only in rural, or in both urban and rural regions, the average African ancestry proportions are higher in South Carolina than in Mississippi. The proportion of African ancestry in JHS (82%) is similar to that observed in their urban and rural samples from Mississippi (83%), while the proportion of African ancestry in Gullah (91%) is slightly higher than that in their rural samples from South Carolina (88%). We thus infer that the slightly higher mean level of African ancestry in the Gullah might be an effect of their rural sampling, but also arose because of historical sociocultural factors.

### Native American ancestry estimates in Gullah African Americans

Consistent with early mtDNA and Y-chromosome studies of Gullah African Americans (22), we found a small Native American contribution to the African American groups sampled for our study. We observed that the Gullah and JHS African Americans had slightly higher Native American ancestry than (~1.3-1.4%) had been reported in most African American groups in the U.S. (2, 3, 5, 6, 8). This discrepancy may reflect the fact that previous studies have often used clustering methods like ADMIXTURE for estimating Native American ancestry proportions which are expected to give underestimates when the proxy population used for Native American ancestry (typically Mesoamerican) is highly genetically drifted from the true source population (Southeastern U.S. Native American). In contrast, the qpAdm ancestry estimation procedure explicitly accounts for genetic drift between the source population and the proxy population and produces an unbiased estimate. In the U.S., only African Americans living in the Southwest U.S. (ASW) from the 1000 Genomes Project had higher Native American ancestry (3.1%) (51).

This level of Native American ancestry is consistent with historical records about the Native American slave trade (18, 52). In the early days of the American colonies, marriages were permitted between Europeans, Africans, and Native Americans (18). Despite a 1671 law forbidding Native American slavery, Native Americans were publicly sold as slaves in Charleston, and their enslavement by colonists was common until the African slave trade accelerated in the 18^th^ century. In fact, from 1670 to 1720, more Native Americans were shipped out of Charleston than Africans were imported (52). After the Yamasee War (1715-17), Native American populations (Yamasee, Ochese, Waxhaw, Santee) declined in South Carolina, and most of the remaining Native American slaves were apparently absorbed into the African community (18). The offspring of Africans and Native Americans were called *mustizoes* or *mustees*, in contrast to the *mulattoes* resulting from the union of Africans and Europeans.

From the 1730s through the 1780s, newspaper ads proclaimed that 2,424 slaves had run away from their masters in South Carolina. Skin color, as noted in 27% of the ads, revealed that a surprising 37% of runaways were light, yellow, or mulatto in appearance, and 19% of them were said to be mustees (18). Cultural influences of Native Americans on the Gullah are reflected in crafts, colono-ware, boat building techniques, or the decoctions of healing herbs used to cope with illness (18). Further support for Native American admixture in the South comes from the several socially distinct communities with European, African, and American Indian ancestry that have persisted to the present day (e.g., Brass Ankles and Turks in South Carolina). In summary, historical, ethnographic, and mtDNA and Y-chromosome data support a Native American contribution to the Gullah (18, 22, 52, 53), which we confirm with our genome-wide data.

### Sex-biased admixture in Gullah African Americans

We found evidence for patterns of sex-biased gene flow in the Gullah (Table 1), consistent with the reported higher male European and female African contributions in other U.S. African Americans (2-6, 8), as well as previous work with the Gullah (22). The significant decrease in European ancestry on the X-chromosome implies a male European ancestry bias, which is consistent with the rape and/or coerced sexual interactions that occurred between European males and African females (54, 55), as well as an overrepresentation of males among African slaves brought to North America (about 70%) (56-59) (e.g., see *Trans-Atlantic Slave Trade Database* website(60)). Notably, these results show that Gullah and JHS African Americans have differing degrees of sex-biased ancestry contributions, with the Gullah exhibiting a greater male-biased European contribution, as shown by a higher proportion of European ancestry coming from males (Table 1). These results are consistent with the negligible European female and higher European male contributions previously noted in the Gullah (22).

As recently reported (8), the extent of this sex bias toward European male and African female genetic contributions is known to vary across the Americas due to regional differences in slavery practices. Despite the lack of direct ethnographic records on mating patterns for the Gullah, this evidence that virtually all European X-chromosomes came from men suggests that few, if any European women married into the Gullah community. However, given the complex mechanistic models of historical admixture (61), any potential scenario that might explain our results is, without historical data, speculative.

### African ancestry of Gullah African Americans

We next tried to elucidate whether the genetic data supported a postulated Sierra Leone (16) or diverse African ancestry (18) for the Gullah. Ancestry estimates (Figure 2) suggested that, relative to JHS African Americans, the Gullah had comparable Yoruba ancestry, and higher ancestry from the African Rice Coast (from Senegal down to Liberia). As shown in Figure 3, a gradient in the clustering of the Gullah and non-Gullah African Americans indicated the Gullah’s relative proximity to the Sierra Leone (especially Creole) and Mandenka samples, while non-Gullah African Americans’ appeared closer to the Yoruba (Figure 3). A quadratic model fit to the regression lines on PC1 and PC2 for the Gullah and non-Gullah African American groups revealed different y-intercepts and slopes for each African American group with a significant interaction term (p = 2.2×10^−16^) (Figure 3). Thus, although not forming individual clusters, the Gullah and non-Gullah samples were distributed along a gradient in this PCA, with the Gullah samples showing more Sierra Leone relatedness than the JHS samples.

**Figure 3.**
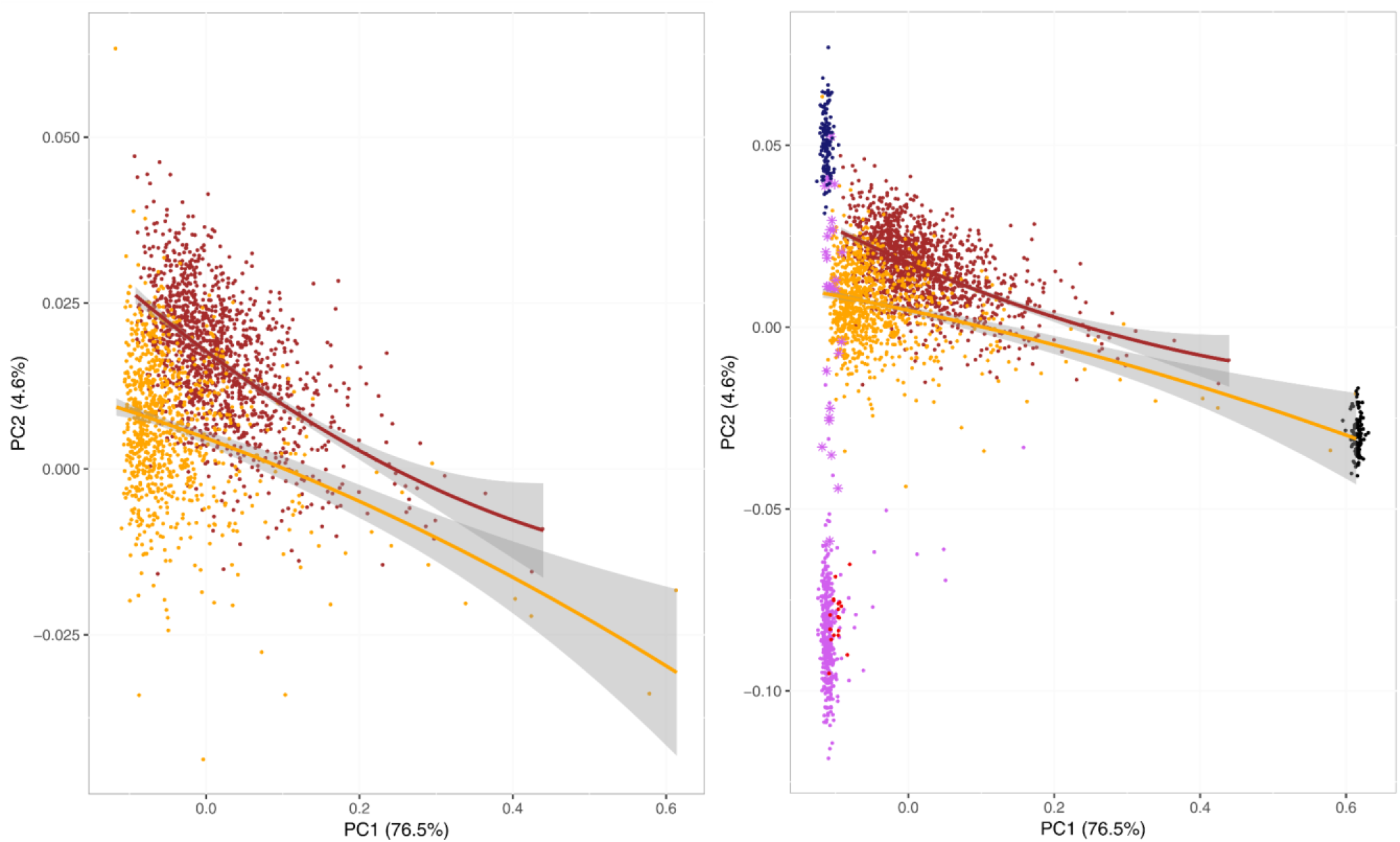
Principal component analysis of Gullah and non-Gullah African Americans. Principal component analysis (EIGENSOFT) using European (CEU samples from Utah) and African ancestral reference populations (YRI samples from Nigeria, Mandenka from Senegal, and Sierra Leone samples) illustrates the Gullah’s closeness to Sierra Leone populations and the Mandenka, and also non-Gullah (JHS) African Americans’ proximity to the Yoruba. To demonstrate the difference between Gullah and JHS African Americans, a quadratic model was fit to the regression lines on PC’ and PC2 for each of the two groups. The resulting interaction term was significant (2.2×10^−16^), indicating a significant difference in the slopes of the two lines. The panel on the left shows the same comparison of Gullah and JHS regression lines without the ancestral population samples. AA: African American; JHS: Jackson Heart Study African Americans from Jackson, Mississippi.

Close relatives are expected to share large identical-by-descent (IBD) segments, which can then be used to model recent ancestry and elucidate population-level relatedness. Analysis of the mean number of shared IBD segments between pairs of Gullah and non-Gullah (JHS) African American individuals confirmed that, relative to Southeast non-Gullah African Americans, Gullah individuals had a lower mean number of shared European segments and a higher number of shared African segments, including a slightly higher proportion of segments of Mandenka ancestry (Supporting Figure S6).

Furthermore, the fixation index (F_ST_) estimates (43) computed to quantify the genetic differentiation between populations, were smaller between the Gullah and the Yoruba and Sierra Leone populations than between the JHS African Americans and these same African populations (Supporting Table S3). The closeness to African populations parallels the higher African ancestry of the Gullah relative to the JHS African Americans. For both Gullah and non-Gullah African Americans, F_ST_ estimates for either Sierra Leone or Yoruba were similar, confirming the similar closeness of each African American group to both African populations. Collectively, these data support the closeness of the Gullah to putative ancestral African populations and the view that the Gullah are not direct and exclusive descendants of populations from Sierra Leone. Instead, as postulated by Pollitzer (18), the Gullah share a common ancestry with numerous populations from Sierra Leone and other regions of West Africa.

These results are consistent with the recently reported genetic ancestries of African Americans from the Southeast U.S. (8). In this large and representative cohort, Micheletti and colleagues found that Southeast African Americans had the highest African ancestry from Nigeria (26-30%), followed by Coastal West Africa (Sierra Leone and the Windward Coast, (~18%)), West Central Africa (~8%), and Senegambia (~7%). There are obvious discordances with the proportions of Africans that arrived in Charleston through the legal slave trade, who were mostly from West Central Africa (~39%), the Windward Coast and Sierra Leone (~23%), and Senegambia (~20%), and only ~5% from the Bights of Benin and Biafra (18). As summarized, the overrepresentation of Nigerian ancestry can be explained by the trade of enslaved people from the British Caribbean (8), and the ethnic composition of Africans imported into the British West Indies indicating source areas in the Gold Coast and the Bights of Benin and Biafra (18). On the other hand, the underrepresentation of Senegambian ancestry can be explained by accounts of early trading and high mortality from this region (8).

### Effects of geographic isolation on Gullah African Americans

Since the Gullah have remained a relatively isolated group over the past few centuries, we sought to determine whether this isolation has affected the genetic structure of their populations. Hallmarks of isolated populations include increased frequencies of recessive disorders, reduced genetic diversity, and higher identity-by-descent (IBD) as the result of founder events and population bottlenecks. There are no reports of the increased frequency of any recessive disorders in the Gullah that would support the occurrence of founder events. We first compared the genetic diversity of the Gullah and Sierra Leone populations by measuring mean heterozygosity and inbreeding coefficients. We observed very similar, though slightly lower level of heterozygosity (P=2.72×10^−3^) (Table 2 and Supporting Table S4) and higher inbreeding coefficient (P= 8.78×10^−3^) (Table 2 and Supporting Table S5, Figure S7), in Gullah compared with Sierra Leone individuals. The similarity in heterozygosity noted in Gullah and Sierra Leone individuals, despite the Gullah’s admixture with European individuals, is not unexpected given their low levels of European admixture. This is because the genetic divergence between two West African chromosomes is similar to that between a West African and a European chromosome.

**Table 2.** Genetic diversity in Gullah African American and Sierra Leone populations

| Population | HETexp | HETobs | *F* |
| --- | --- | --- | --- |
| **Gullah** | 0.332 | 0.333* | -0.0018† |
| **Sierra Leone** | 0.332 | 0.334 | -0.0045 |

Expected heterozygosity (HETexp), observed heterozygosity (HETobs), and inbreeding coefficient (*F*) for the Gullah African American and Sierra Leone populations. † $P < 0.01$ compared with the Sierra Leone population (Wilcoxon test).

A comparison of the different proportions of IBD segments shared in the Gullah and the JHS African Americans showed a lower mean number of shared European segments and a higher number of shared African segments in the Gullah (Supporting Figure S6). This increased number of long founding African haplotypes in the Gullah supports their increased proximity to West African populations, and is consistent with their relative geographic isolation (18). We then used IBDNe, a method based on IBD, to predict the African ancestry-specific effective population size histories for the African American samples over recent generations (42) (Figure 4). The estimated effective population sizes, based only on the African ancestry-associated IBD segments, in African American groups were mostly similar to each other (Figure 4). This result suggested historical mixing within the larger African ancestry population that encompasses these groups, causing the two Southeast African American groups to have a shared demographic history.

**Figure 4.**
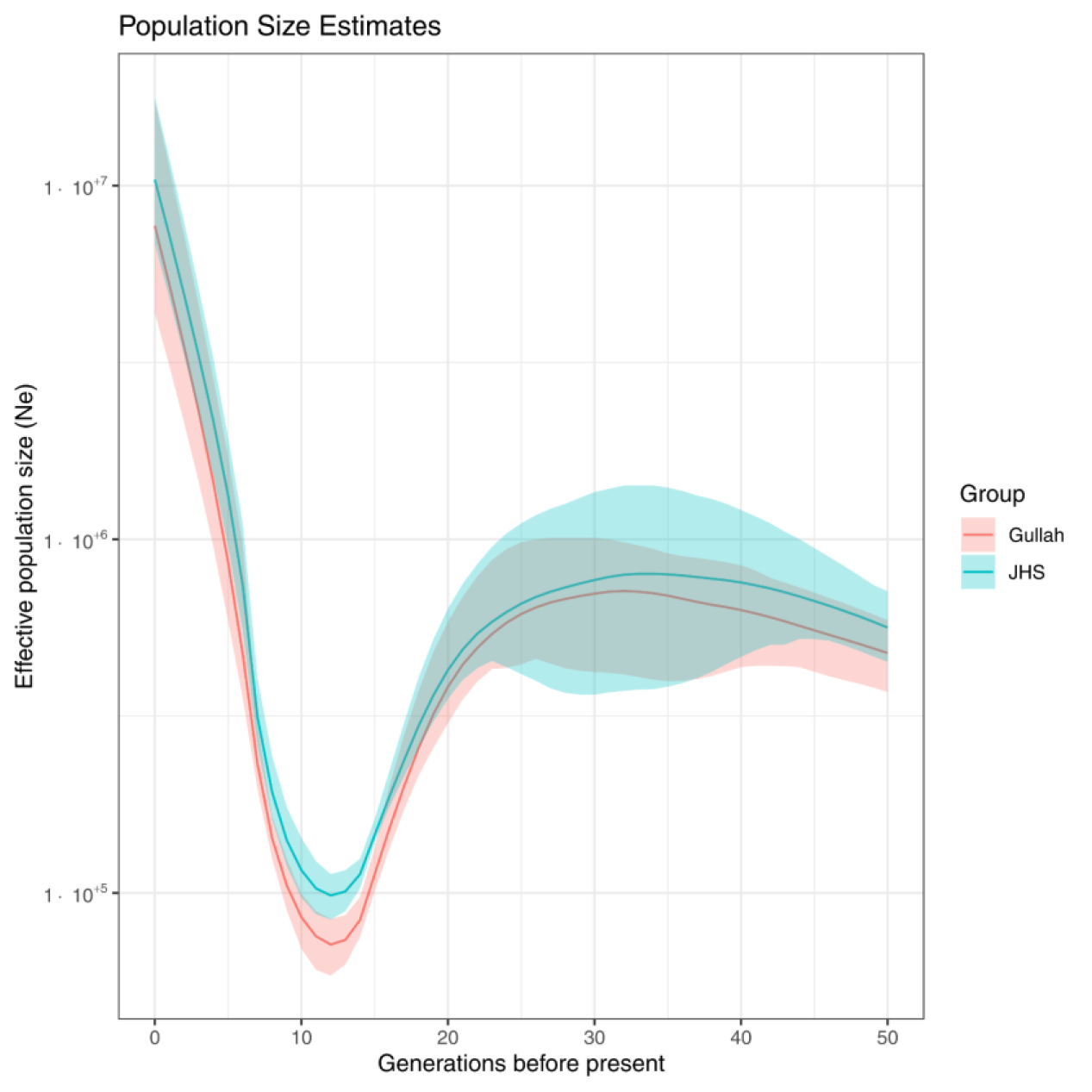
Estimation of ancestry-specific recent effective population size from segments of identity by descent (IBD) in Gullah and JHS African Americans. Plot displays the recent effective population size (Ne) in Gullah and JHS African Americans over the past 50 generations. The lines show the estimated effective population size based on IBD segments associated only with African ancestry, while the colored regions show 95% bootstrap confidence intervals. The graph displays a bottleneck event occurring nearly 13 generations ago, an estimate consistent with the turn of the 18^th^ century. We used GERMLINE to compute sharing of long IBD segments (*l* ≥ 18cM), which are informative of recent relatedness, and used the ancestry-specific IBDNe pipeline to estimate the effective population sizes.

This analysis further revealed a bottleneck event 13 generations ago for both groups (Figure 4), an estimate consistent with the turn of the 18^th^ century. We infer that these bottlenecks mostly resulted from migration and death during the enslavement process. At the same time, these data also support somewhat different demographic histories for the two populations. Based on the African ancestry-associated IBD segments, Gullah have a slightly tighter bottleneck and lower estimated current effective size than JHS, a finding that is consistent with the higher number of long African segments of IBD among Gullah than JHS (Supporting Figure S6). Southeastern non-Gullah African Americans might therefore have a more diverse geographical origin on average than the Gullah, leading to the larger current estimated effective population size. However, the apparently stronger bottleneck in the Gullah could also be a bias due to the reduce European admixture in the Gullah. Collectively, these results are consistent with the relative isolation of the Gullah.

## CONCLUSION

Our study helps clarify the debated ancestry of Gullah African Americans. We confirmed their higher African, lower European, and small Native American ancestries, as well as a larger proportion of male-biased European admixture. We show that the Gullah have a diverse African ancestry, with increased proximity to West African populations than Southeastern non-Gullah African Americans. We found genomic evidence for a slightly tighter bottleneck in the Gullah consistent with a founder event(s) upon importation to the U.S. These findings are consistent with historical, cultural, and anthropological evidence indicating that their relative geographical isolation and strong community life allowed the Gullah to preserve many aspects of their African cultural heritage (18). Although the subtle genetic differences relative to Southeastern non-Gullah African Americans support somewhat different demographic histories, these results also reveal largely shared common ancestries. As such, our data shows that the Gullah are not a genetically distinct group per se, but rather a culturally distinct group of African Americans with subtle variation in its genetic structure.

Broadly, this study shows that subtle differences in genetic structure and ancestry exist. These differences can have important implications for precision medicine, and further reveal the crucial need to include more ancestrally diverse individuals in medical genomic studies (6). Only a comprehensive understanding of the genetic architecture of these populations can ensure that they are not omitted from developments in new genetic technologies and clinical advancements, ultimately contributing to the closure of the health disparities gap as healthcare moves towards precision medicine. Finally, this study is important for Gullah and non-Gullah African Americans who were stripped of ancestral identities by the slave trade. Combined with socio-historical resources, this research can help to recover ancestral histories, and contribute to their new collective identities and ties to ancestral homelands, ultimately paving the road towards transforming lives and possibly reconciliation (13).

## ACKNOWLEDGEMENTS

We thank the Sea Islands Family Project Citizen Advisory Committee (SIFT-CAC) for their enthusiasm and support. We are grateful to the participants from Projects SuGAR, SLEIGH, COBRE and JHS. We thank Yiqi Huang and Kasia Bryc for their contributions to early design, analysis, drafting and revision of this work, as well as Kerry Lok, Xuanlin Hou, Fang Chen, and Yin Lin for technical and analytical assistance. David Reich is an Investigator of the Howard Hughes Medical Institute. This work was supported by National Institutes of Health (NIH) research grants R01 DK084350 (MMS), K01 AR067280 (PSR), P20 RR-017696 and M01 RR001070 (South Carolina COBRE for Oral Health), P60 AR049459 (GSG and DLK), P30 AR072582 (GSG, DLK, PSR), UL1 RR029882 (GSG and DLK), and the W. M. Keck Foundation (WTG).

## DATA AVAILABILITY STATEMENT

All genotypic data used in this study have been deposited in public databases. Gullah African American and African (Sierra Leone) data can be accessed through dbGAP with the accession code phs000433.v1.p1 (The Sea Islands Genetic Network (SIGNET)).

## AUTHOR CONTRIBUTIONS

**Kip D. Zimmerman:** data curation; formal analysis; investigation; validation; visualization; writing-original draft; writing-review and editing.

**Theodore G. Schurr:** investigation; writing-original draft; writing-review and editing.

**Wei-Min Chen:** conceptualization; data curation; investigation; methodology; writing-original draft; writing-review and editing.

**Uma Nayak:** data curation; investigation; writing-original draft; writing-review and editing.

**Josyf C. Mychaleckyj:** investigation; writing-original draft; writing-review and editing.

**Queen Quet:** investigation; writing-original draft; writing-review and editing.

**Lee H. Moultrie:** investigation; writing-original draft; writing-review and editing.

**Jasmin Divers:** data curation; investigation; writing-original draft; writing-review and editing.

**Keith L. Keene:** investigation; writing-original draft; writing-review and editing.

**Diane L. Kamen:** funding acquisition; resources; writing-original draft; writing-review and editing.

**Gary S. Gilkeson:** funding acquisition; resources; writing-original draft; writing-review and editing.

**Kelly J. Hunt:** investigation; resources; writing-original draft; writing-review and editing.

**Ida J. Spruill:** investigation; resources; writing-original draft; writing-review and editing.

**Jyotika K. Fernandes:** funding acquisition; resources; writing-original draft; writing-review and editing.

**Melinda C. Aldrich:** investigation; validation; writing-original draft; writing-review and editing.

**David Reich:** conceptualization; formal analysis; investigation; methodology; writing-original draft; writing-review and editing.

**W. Timothy Garvey:** conceptualization; funding acquisition; investigation; resources; validation; writing-original draft; writing-review and editing.

**Carl D. Langefeld:** conceptualization; formal analysis; investigation; methodology; validation; writing-original draft; writing-review and editing.

**Michèle M. Sale:** conceptualization; funding acquisition; investigation; project administration; resources; supervision; writing-original draft; writing-review and editing.

**Paula S. Ramos:** conceptualization; funding acquisition; investigation; project administration; supervision; validation; writing-original draft; writing-review and editing.

