## Supporting Information; Supporting Figure; Supporting Table. for "Genetic Landscape of Gullah African Americans"

**This PDF file includes:**

Supporting Information Text
Detailed Description of Subjects and Methods
Figures S1 to S7
Tables S1 to S5
References for Supporting Information reference citations

### Supporting Information Text

#### Community engagement

It is important to note that this study was conducted in cooperation with and approval from the Sea Island Families Project (SIFP) Citizen Advisory Committee (1). Interdisciplinary research teams from the Medical University of South Carolina (MUSC) developed community-engaged research projects between the academic researchers and Gullah African Americans residing in rural South Carolina, leading to the formation of SIFP Citizen Advisory Committee (1). This partnership has been ongoing for over 20 years. Participants from three SIFP projects were included in our study: the Sea Island Genetic African American Registry (Project SuGAR) (2); the Center of Biomedical Research Excellence (COBRE) for Oral Health pilot project, "An Epidemiological Study of Periodontal Disease and Diabetes: Cytokine Genes and Inflammation Factors" (3); and the Systemic Lupus Erythematosus in Gullah Health (SLEIGH) Study (4). The SIFP Citizen Advisory Committee meets quarterly for sharing of research results and providing guidance and recommendations to new research. The SIFP Citizen Advisory Committee agreed to this study aimed at understanding the population genetics and ancestry of the Gullah.

### Detailed Description of Subjects and Methods

#### Sample collection and SNP data generation

Self-identified Gullah African American subjects in our study from the aforementioned projects SuGAR (2), COBRE for Oral Health (3), and SLEIGH (4) were recruited under ongoing protocols approved by the MUSC Institutional Review Board and adhered to the tenets of the Declaration of Helsinki. All self-identified Gullah African American participants and their parents were born and raised in the Sea Islands region of South Carolina, or South Carolina low country (along the coastal border and 30 miles inland). The vast majority of Gullah participants were recruited through scheduled community health fairs, recruitment events at local churches, medical clinics, and established organizations on the Sea Islands (2, 5). All subjects received a general medical examination, donated blood samples for DNA analysis, and provided basic demographic and ethnic information. DNA was extracted from blood using a standardized DNA isolation kit (Gentra Systems, Minneapolis, MN). Sample collection and processing have been previously described for the African American subjects from the Jackson Heart Study (JHS) (6, 7), Native American Mixtec (8), and Sierra Leonean (9) subjects. We note that, while Gullah African Americans represent a predominantly rural sample, JHS participants represent the urban, metropolitan area of Jackson, Mississippi. Of note, while the Creole from Sierra Leone were largely recruited in Freetown, and are thus a more urban group, the other ethnic groups were recruited in smaller towns and communities where ethnic communities had lived for many years and the ethnic affiliations were distinct and reliable, and are thus representative of more rural areas. In summary, demographic representativeness was attempted during sampling.

DNAs from the Gullah and Sierra Leone African subjects were genotyped using the Affymetrix Genome-Wide Human SNP Array 6.0 at the Children's Hospital of Philadelphia's (CHOP) Center for Applied Genomics. The Bayesian robust linear model with Mahalanobis (BRLMM-P) algorithm was used to generate SNP calls, and additional quality control (QC) was performed to exclude SNPs with low genotype call rates (<95 %), low minor allele frequency (MAF < 0.05), and genotypes inconsistent with Hardy Weinberg Equilibrium (HWE) ($P < 10^{-10}$). Samples were excluded for low call rate (<95%) and for gender inconsistencies between recorded and genotype-inferred sex. Duplicates, first-, second-, and third-degree relatives were also excluded, with unrelatedness being defined as pair-wise kinship coefficients smaller than 0.0442 estimated by Kinship-based Inference for GWAS (KING) software v2.1.2 (10). Prior to QC procedures, 1,558 Gullah African Americans, 1,775 JHS African Americans, 400 Sierra Leone Africans, and 8 Mixtec individuals were available for our analysis. The Mexican Mixtec individuals have >99% Native American ancestry estimated by ADMIXTURE (11). After QC procedures, 883 unrelated Gullah African Americans, 1,322 unrelated JHS African Americans, 381 unrelated Sierra Leone Africans, and 7 Mixtec subjects were retained for analyses (Supporting Table S1). In addition, 125 unrelated Stanford-Human

Genome Diversity Project (HGDP) (12) and 386 unrelated HapMap III (release 3) (13) subjects were used for analysis. The geographic and linguistic distribution of the African populations used in this study are shown in Supporting Table S2 and Figure 1A.

#### Data merging and SNP trimming

PLINK v1.9 (14) was used to combine our Affymetrix 6.0 data, HapMap III release 3 data, and HGDP data. After merging samples, 136,878 common variant SNPs were retained after removing SNPs: 1) with strand problems, 2) SNPs in the HLA region, 3) MAF<0.01, 4) HWE<$1x10^{-10}$, and 5) missing rate > 0.05. For methods that required a set of linkage disequilibrium (LD)-pruned SNPs (see below), we further removed SNPs with an $r^2 > 0.1$, leaving 64,303 uncorrelated SNPs for analysis. For more specific analyses, identical merging and filtering methods were used to combine the Affymetrix 6.0 data with the HapMap III data. This step resulted in a combined set of 27,374 pruned and filtered SNPs for the Gullah and 29,277 for the non-Gullah (JHS) African Americans.

#### Principal component analysis for inference of population structure

Principal component analysis (PCA) as implemented in EIGENSOFT v6.0.1 (15) was computed with the Sierra Leone African samples combined with all HGDP and HapMap III African samples, for African subpopulation structure inference. For African American population structure inference, HGDP, and HapMap III European samples were added to the African samples as a reference. PCA analyses were performed using the set of LD-pruned 64,303 SNPs described above.

#### Global estimates of admixture

Unsupervised clustering as implemented in ADMIXTURE v1.3.0 (11) was used to estimate global genetic ancestry of the European, African, and African-American populations. ADMIXTURE analysis was performed in European, African (including Sierra Leonean tribes), and African American individuals, assuming 2 through 8 ancestral genetic clusters (*k*=2 through *k*=8) to determine the optimal number of ancestral reference groups. Five clusters (i.e., *k*=5) gave the lowest cross-validation error of *k*=2-8 (Supporting Figure S7). To help order the populations according to their genetic similarities, we used average linkage hierarchical cluster analysis based on the means of each of the five ancestral populations computed by ADMIXTURE v1.3.0 (11) and inter-population similarity matrix of Euclidean distances.

#### Inference of ancestry proportions

To generate estimates of ancestry on the autosomes and chromosome X for both the Gullah and non-Gullah (JHS) African-Americans, we ran qpAdm (16). This program leverages allele frequency correlations between the admixed and source populations with distant outgroups to eliminate potential biases due to genetic drift between the true source populations and the ones used as surrogates for them. The HapMap CEU, HapMap YRI, and Mixtec (kindly provided by Drs. Raghavan and Willerslev (8)) were used as source reference populations for European, African, and Native American ancestry, respectively. It is important to note that, unlike other admixed Native Americans, the northern Amerindian Mixtec individuals (8) have >99% Native American ancestry estimated by ADMIXTURE (11). For the outgroups, the Luhya (LWK), Maasai (MKK), Han Chinese (CHB), Japanese (JPT), Gujarati Indians (GIH), and Toscani (TSI) populations from HapMap were used. qpAdm (16) was run on 798 females from the JHS cohort and 680 females from the Gullah cohort. 523,638 SNPs were used to estimate ancestry on the autosomes and 20,879 SNPs were used to estimate ancestry for the X-chromosome. To determine sex-biased admixture in the African American populations, we examined the African, European and Native American ancestry proportions between the X-chromosomes and the autosomes in both the Gullah and the non-Gullah (JHS) African Americans for equality.

**Detection of genomic segments shared identical-by-descent (IBD) between African American groups and estimation of effective population sizes**

We used GERMLINE v1.5.1 (17) to infer IBD tracts of length 18 cM or longer shared between Gullah and non-Gullah (JHS) African American individuals. Long IBD segments ($l \geq$ 18cM) are informative of recent relatedness. Segments longer than 5 cM identified by GERMLINE have a negligible number of false positives (18). IBD estimation was performed using the set of LD-pruned 64,303 SNPs described above. IBDNe (19) was implemented to estimate ancestry-specific effective population size excluding IBD segments with a "mincm" length shorter than 6, 80 bootstrap samples, and a maximum number of generations to estimate of 100.

**Genetic diversity and population differentiation**

To assess the amount of genetic variation within the Gullah and Sierra Leone populations, heterozygosity (HET) and inbreeding coefficients ($F$) were calculated using genotypic data on 273 healthy Gullahs and 381 Sierra Leoneans. After pruning SNPs in high LD ($r^2 > 0.5$) with PLINK v1.9 (14), the remaining 395K shared SNPs between these populations were used to calculate the HET and $F$ statistics. For each individual, HET and $F$ were estimated based on their expected and observed heterozygous calls, with $F$ = (HETexp − HETobs)/(HETexp). The mean HET and mean $F$ were calculated for each population and a two-sample Wilcoxon test was used to test for a difference in HET and $F$ between populations.

We computed the fixation index ($F_{ST}$) (20) between populations to quantify the genetic differentiation between Gullah and non-Gullah African Americans relative to their African and European ancestral populations. Using genotype data from 273 healthy Gullah, JHS African Americans, Sierra Leone, YRI and CEU samples, we computed the Weir and Cockerham's (1984) $F_{ST}$ (20) as implemented in VCFtools v0.1.13 (21). A total of 479K autosomal SNPs were used for this analysis.

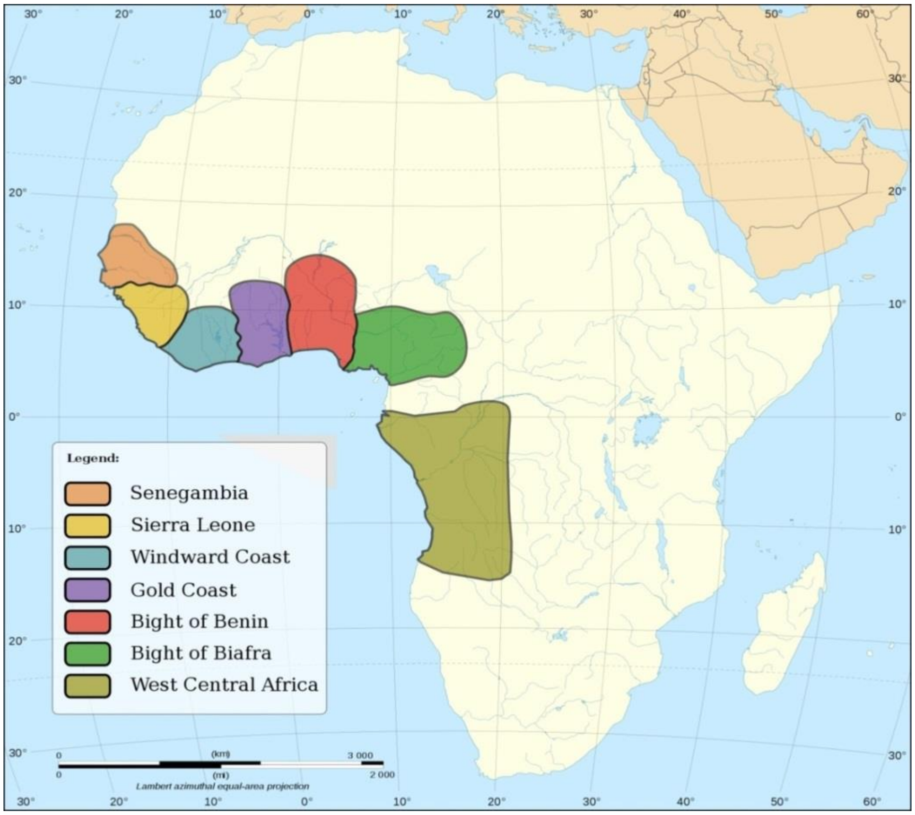


**Figure S1. Major slave exportation regions in Africa during 15th–19th centuries.** Senegambia (Gambia and Senegal), Sierra Leone (Guinea and Sierra Leone), Windward Coast (Ivory Coast and Liberia), Gold Coast (Ghana), Bight of Benin (from the Volta River to the Benin River), Bight of Biafra (east of the Benin River to Gabon), and Angola (west central Africa, including part of Gabon, Congo, and Angola). (Modified by Grin 20 based on the original Africa_map_blank.svg by Eric Gaba under the Creative Commons Attribution-Share Alike 2.5 Generic, 2.0 Generic and 1.0 Generic license and the Creative Commons Attribution-Share Alike 3.0 Unported license. http://en.wikipedia.org/wiki/File:Africa_slave_Regions.svg)

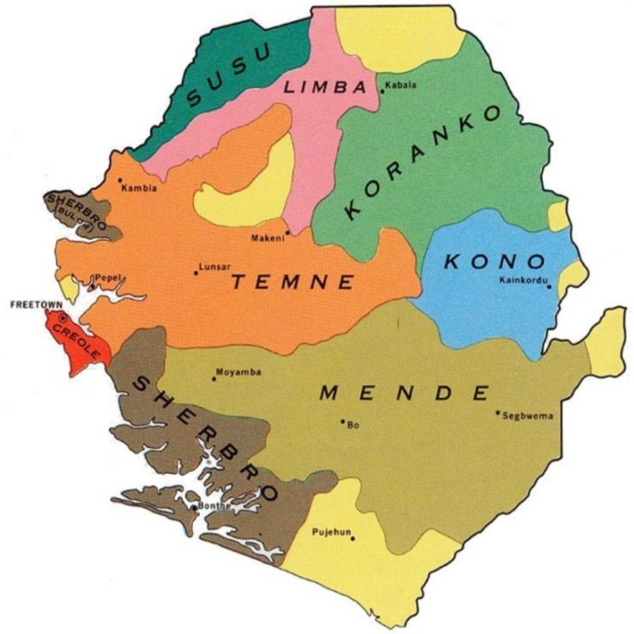


**Figure S2. Sierra Leonean ethnic groups.** Map of Sierra Leone and its ethnic groups groups. Our data set has samples from 10 ethnic groups of Sierra Leone (Creole, Fullah, Kono, Kroo, Limba, Loko, Madingo, Mende, Sherbro, Susu, Temne) (Figure courtesy of the University of Texas Libraries, the University of Texas at Austin).

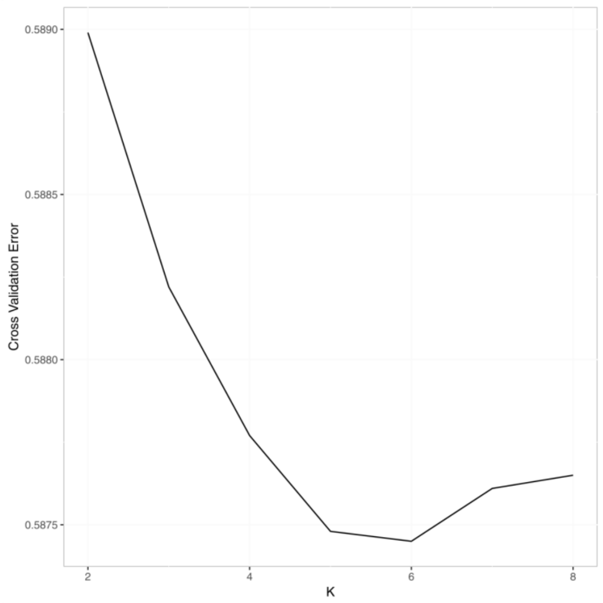


**Figure S3. Cross-validation (CV) error plot in the ADMIXTURE analysis to determine optimal *k* for all subjects.** The most appropriate *k* was 5.

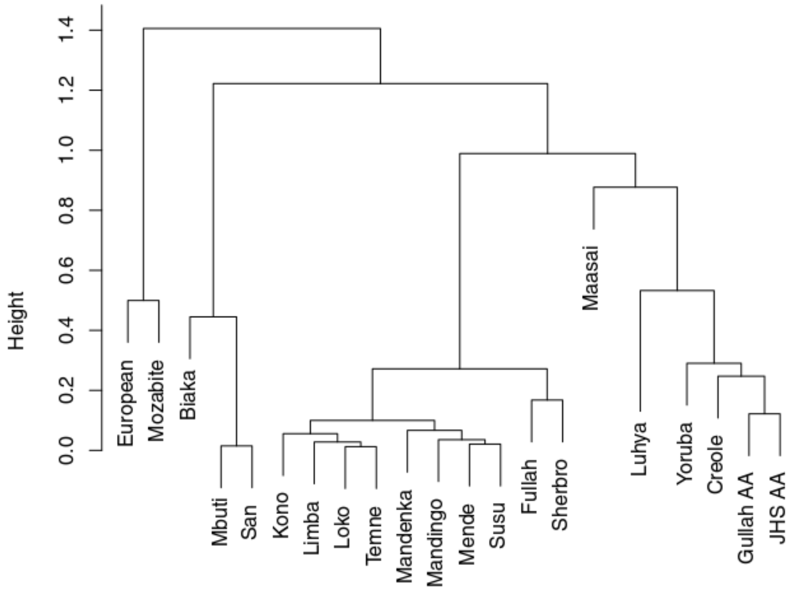


**Figure S4. Dendrogram used to order populations for the ADMIXTURE plot (shown in Figure 2).** Populations were clustered via the means of each the five ancestral populations (*k*). Using the five values for each population, a hierarchical cluster analysis was computed using an inter-population similarity matrix of Euclidean distances. Using the similarity matrix, each population begins as its own cluster and the algorithm proceeds to iteratively join the two most similar clusters until all clusters are joined. AA: African American; JHS: Jackson Heart Study African Americans from Jackson, Mississippi.

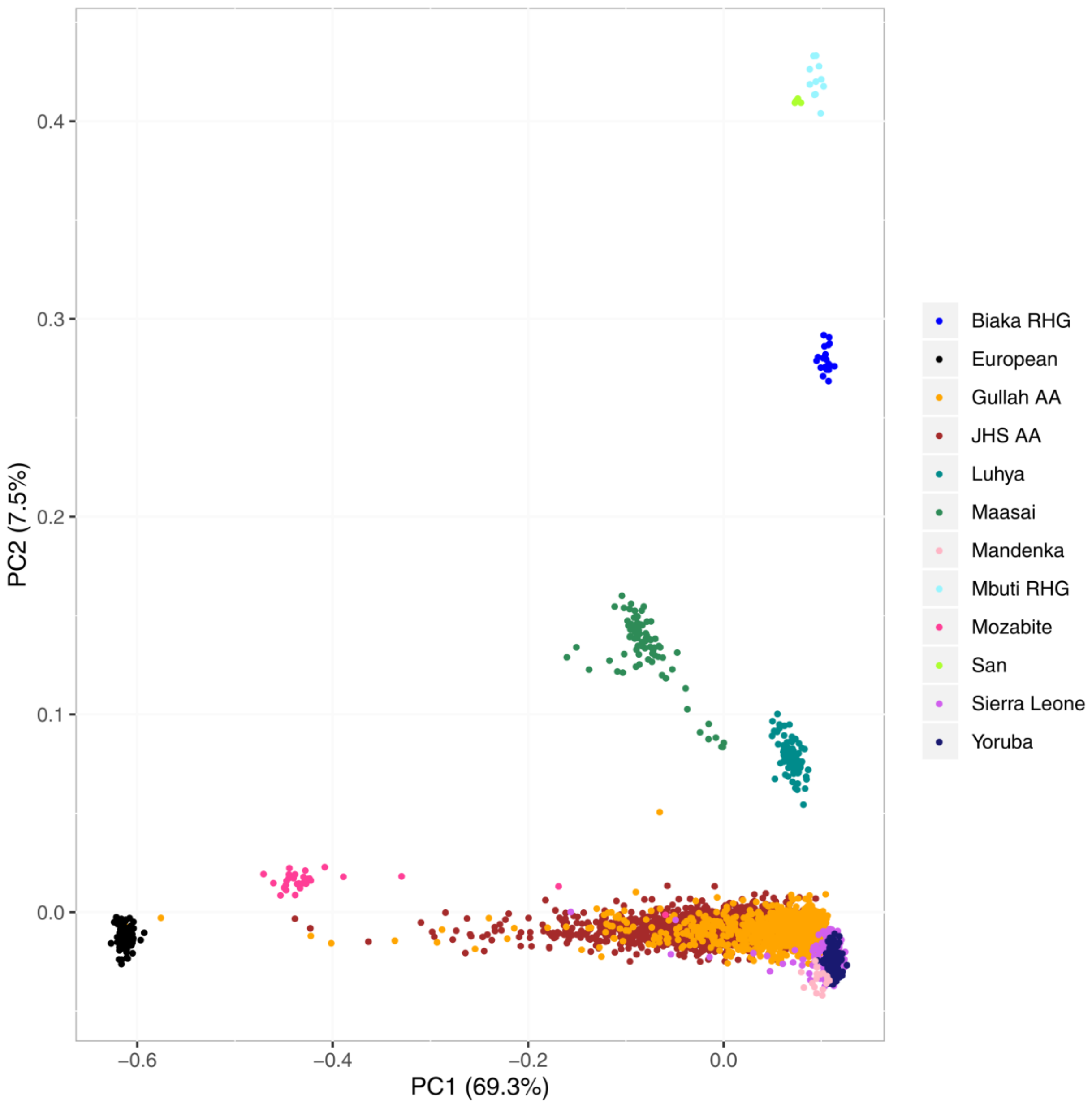


**Figure S5. Principal component analysis of all African and African American samples.** Principal component analysis (PCA) (EIGENSOFT) was applied to HGDP and HapMap III African, African American, and Sierra Leone populations. PCA shows the relative similarity of the African American populations compared to the Sierra Leone, Yoruba, and Mandenka populations.

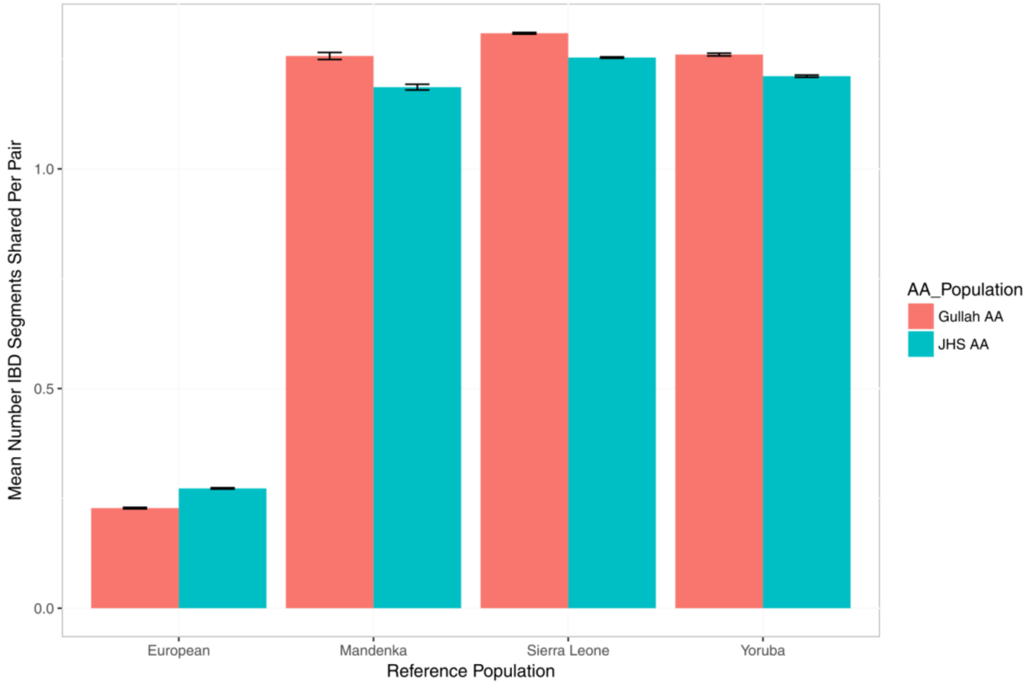


**Figure S6. Identity by Descent (IBD) in Gullah and JHS African Americans reflects their ancestry.** Histogram shows mean number of shared segments of IBD between pairs of individuals. We used GERMLINE to compute the sharing of long IBD segments (*l* ≥ 18cM), which are indicative of recent relatedness. Relative to the JHS, the Gullah African Americans show less European and more African IBD segments. Also, compared to JHS, the Gullah show a slightly higher proportion of segments with the Mandenka.

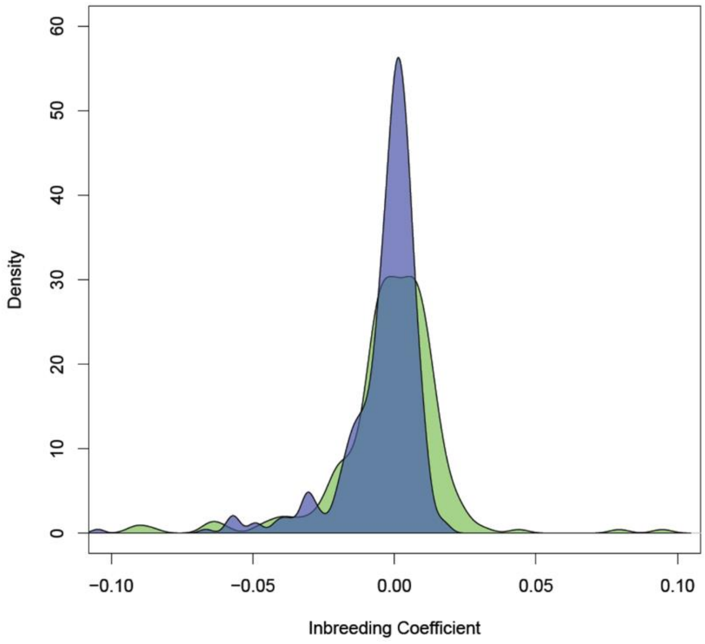


**Figure S7. Density plot of the inbreeding coefficients, *F*, for both the Gullah African American (green) and Sierra Leone (purple) populations.**

**Table S1: African American, Native American, and African Samples and Variants by Study after Quality Control**

| Study | N | Variants | Array |
|---|---|---|---|
| **Gullah AA** | 883 | 699,356 | Affymetrix 6.0 |
| **JHS AA (non-Gullah AA)** | 1,322 | 749,518 | Affymetrix 6.0 |
| **Sierra Leone Africans** | 381 | | |
| Mende | 204 | | |
| Temne | 90 | | |
| Creole | 33 | | |
| Limba | 15 | | |
| Susu | 10 | 699,356 | Affymetrix 6.0 |
| Mandingo | 10 | | |
| Loko | 8 | | |
| Fullah | 6 | | |
| Sherbro | 4 | | |
| Kono | 1 | | |
| **HGDP** | 125 | | |
| Mozabite | 29 | | |
| Biaka RHG | 22 | | |
| Yoruba | 21 | 572,940 | Illumina HumanHap650Y |
| Mandenka | 19 | | |
| Mbuti RHG | 11 | | |
| San | 6 | | |
| **HapMap III** | 386 | | |
| European | 112 | | |
| Yoruba | 111 | 1,201,862 | Affymetrix 6.0 and Illumina 1M |
| Luhya | 80 | | |
| Maasai | 83 | | |
| **Native Americans** | | | Illumina Human660W |
| Mixtec | 7 | 432,378 | |
| **Merged & Filtered** | 3,087 | 136,878 | |
| **Merged, Filtered, & LD Pruned** | 3,087 | 64,303 | |

JHS AA: African Americans (AA) from the Jackson Heart Study (JHS); RHG: rainforest hunter-gatherers.

**Table S2. Geographic and linguistic distribution of the African populations used in this study.**

| Population | Region | Country | Language family |
|---|---|---|---|
| Mozabite | Northern Africa | Algeria | Afroasiatic |
| Yoruba (YRI) | Western Africa | Nigeria | Niger-Kordofanian |
| Mandenka | Western Africa | Senegal | Niger-Kordofanian |
| Sierra Leonean | Western Africa | Sierra Leone | Niger-Kordofanian |
| Biaka RHG | Middle Africa | Central African Republic | Niger-Kordofanian |
| Mbuti RHG | Middle Africa | Democratic Republic of the Congo | Nilo-Saharan |
| Luhya (LWK) | Eastern Africa | Kenya | Niger-Kordofanian |
| Maasai (MKK) | Eastern Africa | Kenya | Nilo-Saharan |
| San | Southern Africa | Namibia | Khoisan |

RHG: rainforest hunter-gatherers. Note: Regions are defined based on the UN Statistics Division geoscheme (https://unstats.un.org/ unsd/methodology/m49).

**Table S3. Pairwise $F_{ST}$ estimated between populations**.

| | Sierra Leone | YRI | CEU |
|---|---|---|---|
| **Gullah African American** | 0.003 | 0.004 | 0.099 |
| **Non-Gullah African American** | 0.038 | 0.038 | 0.109 |

Non-Gullah are African Americans from the Jackson Heart Study.
YRI = Yoruba, and CEU = CEPH European-Americans.

**Table S4. Expected heterozygosity (HETexp) and observed heterozygosity (HETobs) for the Gullah African American and Sierra Leone populations.**

| Population | HETexp | | | HETobs | | | Wilcoxon Test P-value |
|---|---|---|---|---|---|---|---|
| | Mean | Median | SD | Mean | Median | SD | |
| **Gullah** | 0.332 | 0.332 | ±0.00017 | 0.333 | 0.332 | ±0.0064 | 2.73E-03 |
| **Sierra Leone** | 0.332 | 0.332 | ±0.00019 | 0.334 | 0.332 | ±0.0049 | |

SD: standard deviation.

**Table S5. Inbreeding coefficient (*F*) for the Gullah African American and Sierra Leone populations**

| Population | *F* | | | Wilcoxon Test P-value |
|---|---|---|---|---|
| | Mean | Median | SD | |
| **Gullah** | -0.0018 | 0.0005 | ± 0.019 | 8.78E-03 |
| **Sierra Leone** | -0.0045 | -0.0004 | ± 0.015 | |

SD: standard deviation.
